# ENGINEERING AND PRECLINICAL EVALUATION OF DRIED FOOD-FORMULATED YEAST-SECRETED ANTIBODIES AS PERI-EXPOSURE PROPHYLAXIS AGAINST CLOSTRIDIOIDES DIFFICILE DISEASE

**DOI:** 10.1101/2025.11.14.688569

**Authors:** Charlotte Roels, Marlies Ballegeer, Semiramis Yilmaz, Hendrik Grootaert, Leander Meuris, Elise Wyseure, Thomas Herman, Pieter Van Haverbeke, Sandrine Vanmarcke, Katrien Claes, Kenny Roose, Nico Callewaert

**Affiliations:** VIB Center for Medical Biotechnology, VIB, Ghent, Belgium; Department for Biochemistry and Microbiology, Ghent University, Ghent, Belgium

## Abstract

*Clostridioides difficile* gastrointestinal infection (CDI) causes significant disease burden, often following gut dysbiosis induced by antibiotic treatment in at-risk individuals. A non-invasive, easy to use targeted prophylaxis in such cases of acute elevated CDI risk, especially in those with a history of recurrent CDI, would prevent much suffering. In this study, we envisioned a simple prophylactic intervention for CDI enabled by the oral administration of yeast-secreted dried food-formulated *C. difficile* toxin A (TcdA)-and B (TcdB)-neutralizing VHH-based antibodies. We engineered two antibody versions: a previously reported tetravalent VHH (VHH_4_) comprising dual specificities for TcdA and TcdB, and its murine IgA Fc fusion counterpart (VHH_4_-mIgA Fc). We explored Generally Recognized as Safe (GRAS) *Komagataella phaffii* yeast as a low-cost, highly scalable production platform. In a first experiment, we produced the antibodies at small scale in shake flask yeast cultivations and formulated the secreted antibody-containing growth medium into standard-composition dried mouse feed pellets. The antibody food pellet-administration intervention was tested in a *C. difficile* spore-challenged mouse model in which CDI leads to 50% mortality of untreated animals. We started the antibody-feed prophylaxis dosed at 10 nmol/g pellet one day prior to *C. difficile* challenge and continued until the infection was cleared with *ad libitum* feeding. We found that the VHH_4_ feed almost completely prevented mortality and reduced disease-related weight loss. Despite its higher *in vitro* potency against TcdA and equipotent potency against TcdB, the VHH_4_-mIgA Fc fusion did not confer additional *in vivo* benefit, and further work focused on the simpler VHH_4_ format. Scalable high-cell-density bioreactor production of VHH_4_ yielded equally effective prophylactic formulations at the same dose of 10 nmol/g pellet, achieving complete survival and reduced disease burden. Our findings suggest that, with further antibody- and process development, oral prophylaxis with scalable, cost-effective, yeast-produced, dried food-formulated anti-toxin VHHs could offer a simple non-invasive, non-antibiotic prophylaxis option for the high-risk (r)CDI patient population.

## Introduction

The spore-forming pathogen *Clostridioides difficile* (*C. difficile*) most commonly causes nosocomial infection upon antibiotic-induced dysbiosis, which can lead to persistent infections causing toxic megacolon, sepsis and death in severe cases^1,2^. Highly resistant spores are known to cause hard-to-treat, recurrent disease (rCDI) and dangerous and costly hospital or community-acquired outbreaks. Long-term surveillance studies and large meta-analyses indicate that 1-2 % of current clinical strains already show vancomycin resistance^3,4^. Fidaxomicin resistance remains highly uncommon, since its introduction in 2011 in the CDI standard of care (SOC), yet singular cases have also been reported and continued surveillance is needed in the coming decades^5,6^. With up to 15 % of individuals being healthy carriers of *C. difficile*^7,8^, reaching a relative abundance of > 0.1 %, or billions of cells, of the total gut microbiome in CDI patients^9^ and an upsurge of antimicrobial resistant (AMR) strains worldwide, the infection is classified as an urgent threat by the Centers for Disease Control and Prevention (CDC)^3,10–12^. *C. difficile* typically blooms in the context of antibiotics treatment-caused dysbiosis, risk factors include hospitalization after surgery, older age, underlying conditions mainly impacting the immune system, or a previous episode of CDI, forming a clear at-risk target population for currently non-existing preventive therapy^13^. In conclusion, this infection remains a significant public health threat which underscores the continued need for effective preventive and non-antibiotic treatment strategies against CDI and rCDI.

Three well-characterized key virulence factors of *C. difficile* are generally known: large multimeric Toxin A (TcdA, 308 kDa) and Toxin B (TcdB, 270 kDa), both part of the Large Clostridial Toxins (LCT) family, and a smaller enzyme named *C. difficile* binary toxin (CDT, 48 kDa), expressed by only a subset of strains^14,15^. The CDI pathogenesis is largely mediated by the glucosyltransferase activity of the LCTs, with TcdB as the more potent cytotoxin and current evidence in animal models as well as human infection studies strongly suggesting that TcdA and B are involved in most infections but with TcdB more responsible for severe disease^16^. Accordingly, bezlotoxumab, a systemically administered TcdB-neutralizing mAb, is clinically efficacious, but requires disruption of the gut permeability by the *C. difficile* toxins, allowing paracellular transport of the therapeutic antibody from the systemic circulation to the gut lumen^17,18,19,20^. Therefore it is currently indicated only as a concomitant drug to antibiotic SOC in high-risk patients for recurrence, or to treat severe cases of rCDI^21,22^. Unfortunately, Merck has discontinued bezlotoxumab (Zinplava™) sales since January 2025 without clearly citing the reason, though a confluence of limited market uptake due to its narrow patient indication range, uneven reimbursement in different markets, together with an identified safety signal in patients with chronic heart failure may have made it commercially unattractive for the manufacturer to continue this drug.

In previous work from our lab, we demonstrated that a dried feed-admixed formulation method of VHH-based antibodies secreted by *Komagataella phaffii* yeast (commonly known as the *Pichia pastoris* biotech-favored expression system) was very effective in preventing Enterotoxigenic *Escherichia coli* (F4-ETEC) infection in post-weaned minipigs, which are very sensitive for a few weeks due to the interruption of maternal milk antibody supply prior to full development of the piglet’s anti-*E. coli* immunity^23^. This is a major problem in commercial pig farming, requiring a massively scalable low-cost manufacturing and feed-formulation, which is achievable by large-scale *Pichia* bioreactor processes. This product is presently in the final stages of being geared up for large-scale commercial use under the trade name Nanoprotec™^24^. In that work we used VHH-porcine IgA Fc fusions targeting bacterial flagellae. The most likely main proposed mechanism of action was through enchained growth of the bacteria by cross-linking, which prevents the separation of daughter cells as they divide. The infection is then effectively cleared by the natural GI transit, reducing the infection load in the gut<u>^1,2^</u>^,25^. Here, we investigated whether a similar approach, now targeting the *C. difficile* TcdA and TcdB, could be used to prevent CDI, which is a colon infectious disease rather than ETEC, which is a duodenal/ileal disease^13,27^. This anatomical difference presents additional challenges for oral biologic delivery, as therapeutic proteins must remain stable and active throughout the upper gastrointestinal tract before reaching the distal site of infection.

For our exploration, we built on previous work that had designed a tetrameric VHH (VHH_4_) with high multispecific potency against both TcdA and TcdB^28,29^. We engineered *K. phaffii* for constitutive secretory expression of this VHH_4_, as well as the same VHH_4_ C-terminally fused to a murine IgA Fc tail (VHH_4_-mIgA Fc), in line with the anti-ETEC antibody format described above. Sticking rigorously to fully scalable manufacturing unit processes derived from industrial food-processing enzyme manufacturing practice, we admixed the clarified and diafiltration-concentrated anti-toxin antibody-containing fermentates (and control fermentate) with standard mouse feed matrix into doughs, which were extruded, cut and dried as standard mouse feed pellets. We validated that the antibodies were extractable back in functional form from these feeds, and explored their *in vivo* efficacy in the preventive treatment of CDI in the mouse model.

## Results

### *K. phaffii-*produced VHH_4_ and VHH_4_-mIgA Fc antibodies show high *in vitro* toxin-neutralization potencies

To evaluate the therapeutic potential of orally delivered antibodies to prevent CDI, a tetravalent VHH (VHH_4_) was selected from literature^29^ and cloned into *K. phaffii* (also: yeast or *Pichia*), either alone or fused to murine IgA (VHH_4_-mIgA Fc) (**Fig. 1A**). To investigate the quality and potency of *Pichia*-produced VHH(–mIgA Fc) constructs, shake flask (SF) expressions were performed. Upon purification, the HIS-tagged VHH_4_-containing antibodies appeared fully intact when compared to their theoretical and deglycosylated MW when assessed on reducing SDS PAGE after endoH pre-treatment, both via Coomassie BB staining (**Fig. 1B, left**) and western blotting using anti-HIS detection (**Fig. 1B, right**). Productivity in small-scale uncontrolled SF runs increased when switching from P_GAP_ to P_UPP_-driven constitutive expression, with titers reaching 90 mg/L for VHH_4_ and 53 mg/L for the mIgA Fc fusion under P_UPP_ (**Suppl. Fig. 1A)**.

**Figure 1.**
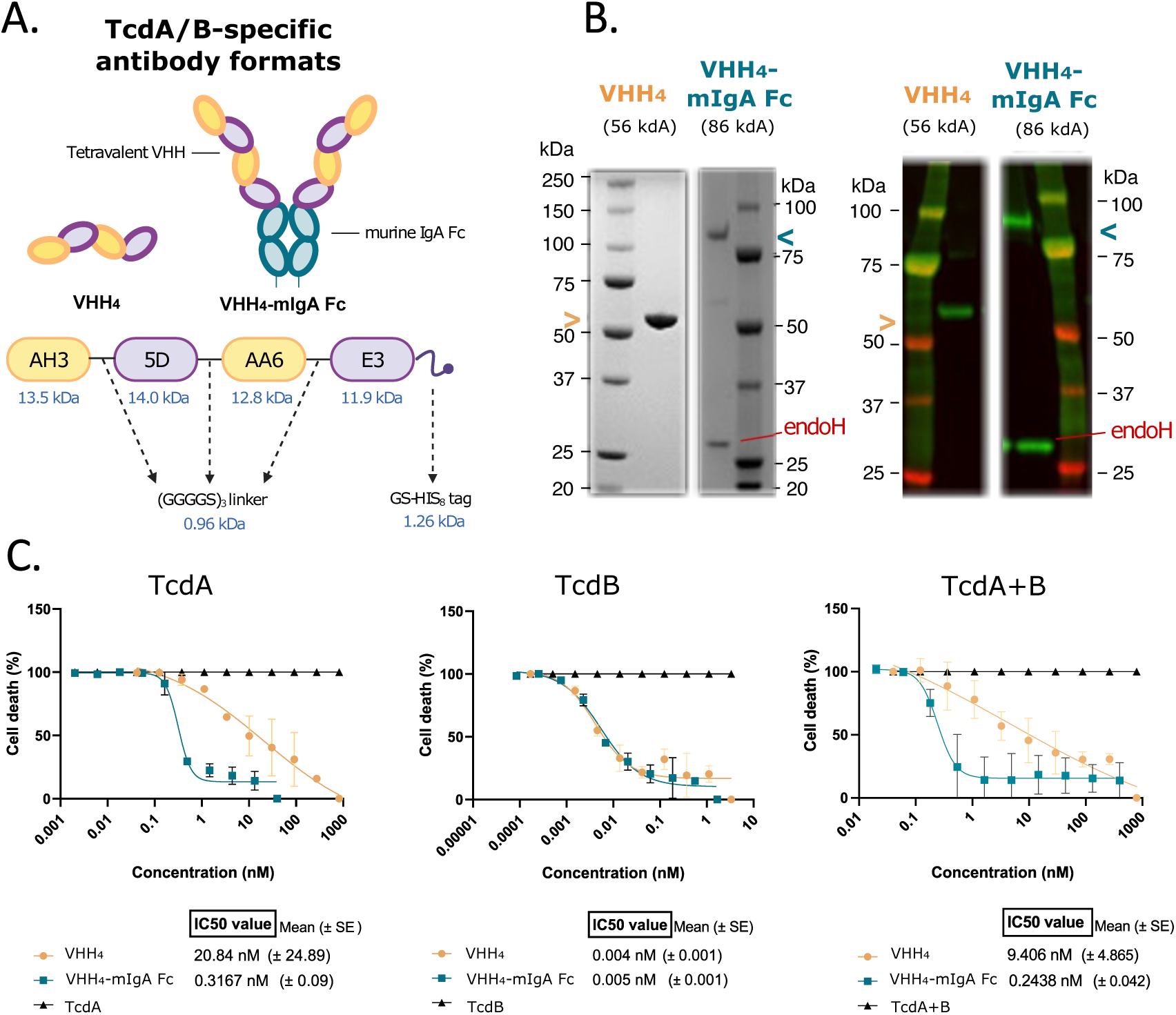
Potent *K. phaffii*-expressed VHH_4_ containing antibodies reach the colon after in-feed oral delivery. (A) Schematic of the tetrameric VHH_4_ and octameric VHH_4_-murine IgA Fc (mIgA Fc) constructs used for expression in *K. phaffii*. Visual created using BioRender.com. (B) Reducing SDS-PAGE (left) and anti-HIS western blot (right) analysis confirms integrity and purity of shake flask-produced VHH4 and EndoH-treated VHH_4_-mIgA antibodies. (C) Vero cell-based luminescence assay showing cell death neutralization potency against TcdA (left), TcdB (middle), and an equipotent toxin mix (right) at low nM potencies, with an almost 50-fold higher potency for VHH4-mIgA over VHH4 against TcdA (n = 3 biological replicates).

To evaluate the neutralization potency of the produced VHH(-mIgA Fc) proteins, the purified anti-toxin antibodies were further characterized by titration against a TcdA/TcdB equipotent mix in a Vero cell-based luminescent reporter assay (**Fig. 1C**). Time-dependent toxin-induced cell death could be observed microscopically (**Suppl. Fig. 1B, left**). Generally, this observation corresponded well with luminescent signal intensity measured in the reporter assay, which also showed 1000-fold higher potencies of TcdB (EC50 = 2-4 pg/mL) over TcdA (EC50 = 2-6 ng/mL) (**Suppl. Fig. 1B, right**). Our lead molecules, VHH_4_ and VHH_4_-mIgA Fc showed high potencies against both TcdA and TcdB alone, and against an equipotent mix of both toxins – to which they exhibit an IC50 value of 9.41 nM (± 4.87 nM) for the VHH_4_ and 0.24 nM (± 0.04 nM) for the VHH_4_-mIgA Fc fusion (**Fig. 1C, right**).

To compare *in vitro* potencies and to determine theoretical pellet dosing of our lead molecules for *in vivo* assays, we defined 1 unit as the amount of anti-toxin antibody that is needed to neutralize 50 % of the *in vitro* cytotoxicity caused by 1 nmol of the pure toxins or 1 nmol of an equipotent mix of both TcdA/TcdB toxins (**Table 1**). For VHH_4_, the potency against TcdB is about 3 times higher than against TcdA. For VHH_4_-mIgA Fc, the potency against TcdB is very similar, yet the multivalency imparted by the fusion of VHH_4_ to the IgA-chain increases the potency against TcdA with a factor of 25. Of note, the observed potency was tremendously lower for the monomeric VHH building blocks as compared to the multimeric constructs, with 0.1–0.4 mU/nmol for anti-TcdA (AH3 and AH3-mIgA Fc) and almost non-existent neutralization capabilities for anti-TcdB (E3 and E3-mIgA Fc) monomers (**Suppl. Fig. 1C**).

**Table 1.**
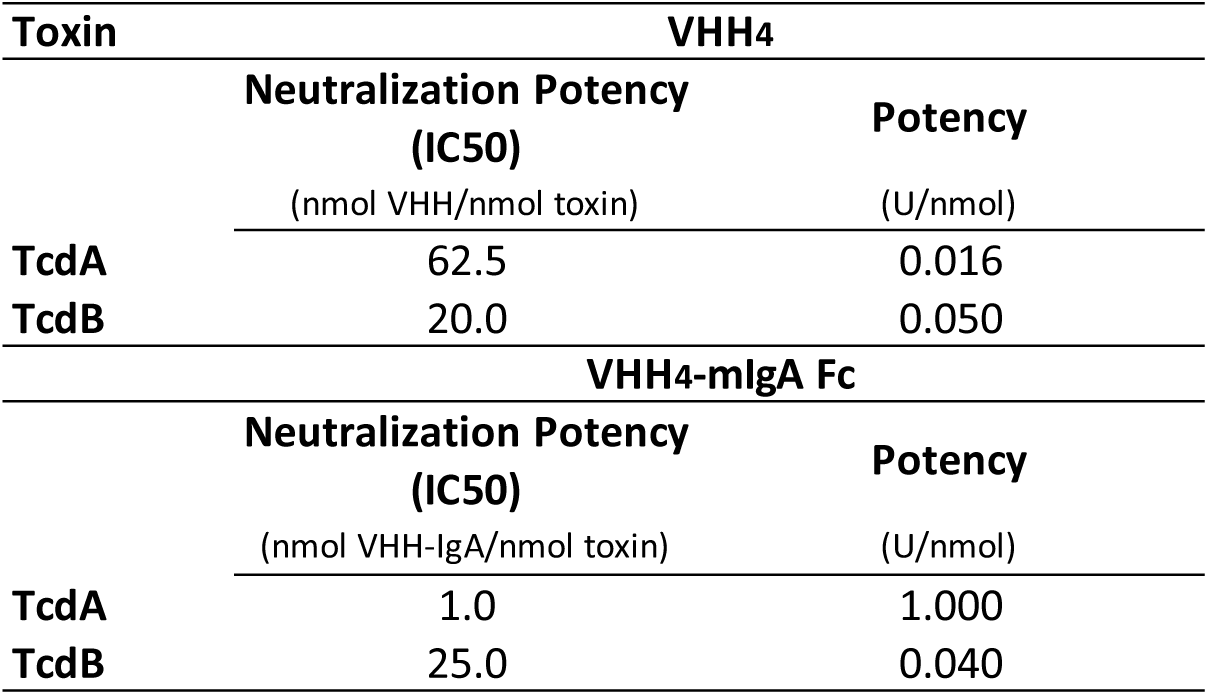
Comparative Neutralization Potency of VHH_4_ and VHH_4_-mIgA Fc against TcdA and TcdB based on *in vitro* Vero cell neutralization assays (n = 3).

| Toxin | VHH<sub>4</sub> |  |
| --- | --- | --- |
|  | Neutralization Potency<br>(IC<sub>50</sub>)<br>(nmol VHH/nmol toxin) | Potency<br>(U/nmol) |
| TcdA | 62.5 | 0.016 |
| TcdB | 20.0 | 0.050 |
| Toxin | VHH<sub>4</sub>-mIgA Fc |  |
|  | Neutralization Potency<br>(IC<sub>50</sub>)<br>(nmol VHH-IgA/nmol toxin) | Potency<br>(U/nmol) |
| TcdA | 1.0 | 1.000 |
| TcdB | 25.0 | 0.040 |

### Production of VHH_4_(-IgA Fc) and formulation in dried mouse feed pellets

To provide the molecules needed for the *in vivo* experiments, both VHH_4_ and VHH_4_-mIgA Fc were obtained from 5 L shake flask (SF) productions. The titer of fully intact protein was evaluated throughout the downstream process steps (DSP) (**Suppl. Fig. 2A**). Dot blots detecting the HIS_8_-tagged molecules were used to follow up product concentration and loss-of-product to the permeate during ultrafiltration/diafiltration (uf/df), showing concentrated levels of VHH_4_ in the retentate sample with little protein loss towards the permeate (**Suppl. Fig. 2B**). HIS-capture ELISA confirmed that little HIS-tagged protein was detected in the permeate at both the uf/df (5 MWCO) and cross flow filtration (CFF, 10 MWCO) step, indicative of little loss of fully intact product during the DSP (**Fig. 2A**). In total, 0.45 g of VHH_4_ and 0.78 g of VHH_4_-mIgA Fc was obtained after DSP, resulting in respective yields of 90 mg/L (1.59 μM) for VHH_4_ and 156 mg/L (0.83 μM) for VHH_4_-mIgA Fc during these larger-scale production runs.

**Figure 2.**
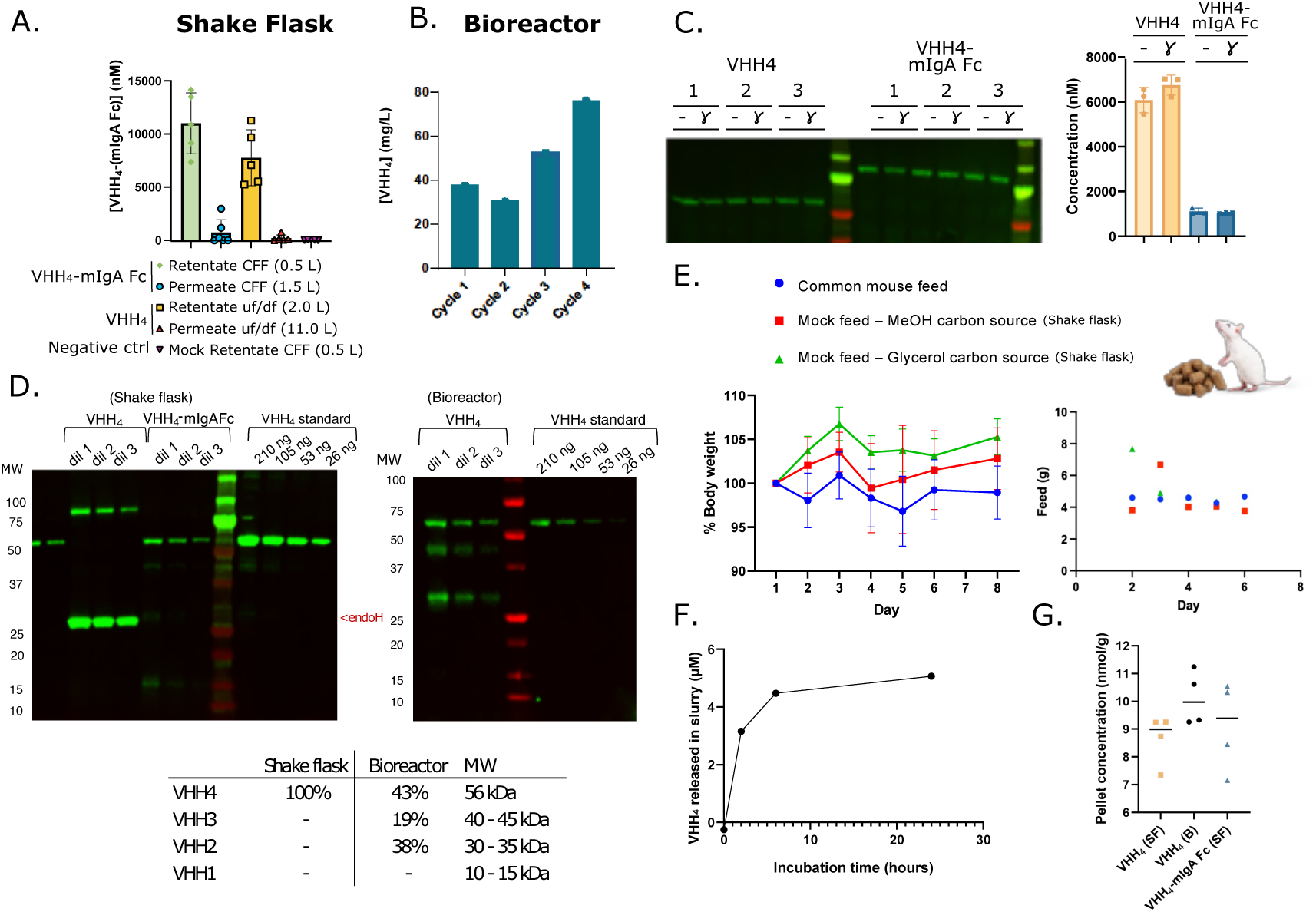
Downstream processing, formulation, and stability of VHH_4_-containing feed pellets. (A) Sandwich ELISA confirms recovery of shake flask-produced antibodies during DSP with little loss to the permeate. (B) Bioreactor-based production of VHH_4_ exhibits increasing yields over repeated fed-batch cycles, with an average yield of 50 mg/L after DSP. (C) Anti-VHH western blot of formulated Exp 1 pellets shows that antibody presence and integrity is preserved during sterilizing γ-irradiation based on both WB (left) and sandwich ELISA (right). (D) Shake flask-produced VHH_4_ appears intact on anti-HIS WB, while bioreactor-produced VHH_4_ shows a fragmentation pattern. Densitometry shows that 100 % vs 43 % of VHH4 is fully intact after pelleting when produced by shake flask or bioreactor runs respectively. (E) Body weight (left) and feed intake (right) data from a palatability study in mice testing yeast HCP-containing mock feed pellets. Two outliers in feed intake occurred on day 1 and 2 post-feed change, indicative of an increased ingestion of pellets during the first days of the experiment (n = 6). (F) *In vitro* VHH4 release kinetics of VHH_4_ from pellet formulation, followed up by ELISA. (G) Quantitative ELISA results show a dose of nearly 10 nmol/g pellet for all experimental pellet batches (n = 4 technical replicates). (H) Investigation of the toxin neutralization potential after 1-2 years of storage confirms retained *in vitro* biological activity present in all POI-containing batches used in *in vivo* experiments (n = 3 biological replicates).

To establish a pre-clinical administration form for edible delivery of yeast-produced antibodies, a lab-scale formulation method was set up, incorporating the lead molecules into mouse feed and generating pellets after extrusion and subsequent air drying at 40 °C for 20 hours (**Suppl. Fig. 2D)**. After formulation, the pellets were sterilized by γ-irradiation (25 kGy over 5 hours), and we validated that extractable quantity and quality of the antibodies was unaltered through this sterilization procedure (using a sandwich ELISA capturing the C-terminal HIS_8_ tag and detecting the alpaca IgG (V_H_) regions of the VHH_4_ (**Fig. 2C, right**), and by anti-HIS-based immunoblot, where band intensity before and after γ-irradiation was not significantly different (**Fig. 2C, left** n = 3 pellets). Drying of the pellets at 30 °C gave a significantly higher microbial load when compared to drying at 40°C or 50 °C. Choosing a condition, we settled on a condition of mixing 700 ml of concentrated yeast culture medium per kilogram of powdered feed, as this resulted in complete moisture removal with reduced microbial load, followed by γ-irradiation of the dried pellets for full sterilization (**Suppl. Fig. 2E**) prior to the *in vivo* experiments.

After formulation of the SF and bioreactor-produced lead molecules into pellets, the quality of the antibodies was further investigated. The absence of clipping of the VHH_4_ (56 kDa) and endoH-treated VHH_4_-mIgA Fc (86 kDa) was confirmed via anti-VHH western blots following extraction of the soluble fraction from the pellet matrix (**Fig. 2D, left**). At this resolution, densitometry indicates that the SF-produced VHH_4_ and VHH_4_-mIgA Fc are both fully intact (**Fig. 2D, table**).

To quantify the extractable amount of VHH_4_(-IgA Fc) in the pellets for the *in vivo* experiments, the preparation of the pellet slurry extract was optimized, and subsequent quantitative ELISA was performed. This effort included the introduction of sieving steps of the powdered feed, and the evaluation of different incubation times and temperatures to accurately assess drug release. We found that VHH_4_ was released quickly from the pellet formulation, when prepared as described in Material and Methods, and reaches a plateau around 6-10 hours (**Fig. 2F**). Based on quantitative ELISA, at this plateau, around 90% recovery is obtained of the 10 nmol/g HIS_8_-tagged VHH_4_-IgA molecules that were formulated into the pellets (**Fig. 2G** and **Table 2**).

**Table 2.** ELISA results showing antibody concentrations in supernatant following 24h drug release from pellet formulation. Mean and standard deviation (SD) of all three experimental pellet batches to be tested in *further in vivo* experiments are depicted (n = 4 technical replicates).

| Pellet concentration - ELISA |  |  |
| --- | --- | --- |
|  | Mean (nmol/g) | SD (nmol/g) |
| VHH<sub>4</sub> (Shake flask) | 8.6 | 0.9 |
| VHH<sub>4</sub>-mIgA Fc (Shake flask) | 9.1 | 3.2 |
| VHH<sub>4</sub> (Bioreactor) | 10.1 | 1.0 |

### Edible peri-exposure prophylactic treatment with a daily dose of 10 nmol/g of VHH_4_ in dried feed pellets shows significant efficacy in a preclinical spore-challenge model of CDI, whereas VHH_4_-mIgA Fc does not

To investigate the efficacy of orally administered anti-toxin VHH_4_(-mIgA Fc) against CDI, we have chosen to use the widely adopted murine antibiotic cocktail-induced dysbiosis model^30^. This model has been described to mimic several key features of CDI in humans, inducing diarrhea, weight loss and histological damage to the colon. After pre-treatment of the mice with a broad-spectrum antibiotics cocktail through the drinking water to eradicate the healthy microbiota (**Suppl. Fig. 3A**), mice develop CDI after *in situ* germination of spores administered via oral gavage (**Fig. 3A)**. In addition, the severity of the disease can be varied by changing the number of spores used for the challenge, or by using different toxicogenic strains. Two ribotypes (RT) of *C. difficile*, RT078 and RT012, were tested for sporulation efficacy. The sporulation efficiency of both strains was in the same range and about 5E8 spores were obtained from 10 Petri plates (**Suppl. Fig. 3B**). We chose to work with RT078 for *in vivo* studies, as this is an emerging strain mostly prevalent in Europe and associated with more severe disease outcomes. Moreover, it was the second most prevalent strain in Belgian hospitals in 2019^31–33^. Spore titration experiments in C57BL/6 mice revealed variable disease progression at lower inoculum levels, whereas a challenge dose of 10⁵ spores/mouse reproducibly induced acute, fulminant colitis, as evidenced by pronounced body weight loss and a mortality rate of 80–90 %, which is excessive and precludes an accurate measurement of any impact of a peri-exposure prophylactic intervention on the post-acute recovery dynamics. Consequently, a smaller inoculum with 10^4^ spores/mouse was chosen. This resulted in a somewhat more moderate disease course, but still resulting in 30–50 % mortality in the untreated control cohort (**Suppl. Fig. 3C-D**).

**Figure 3.**
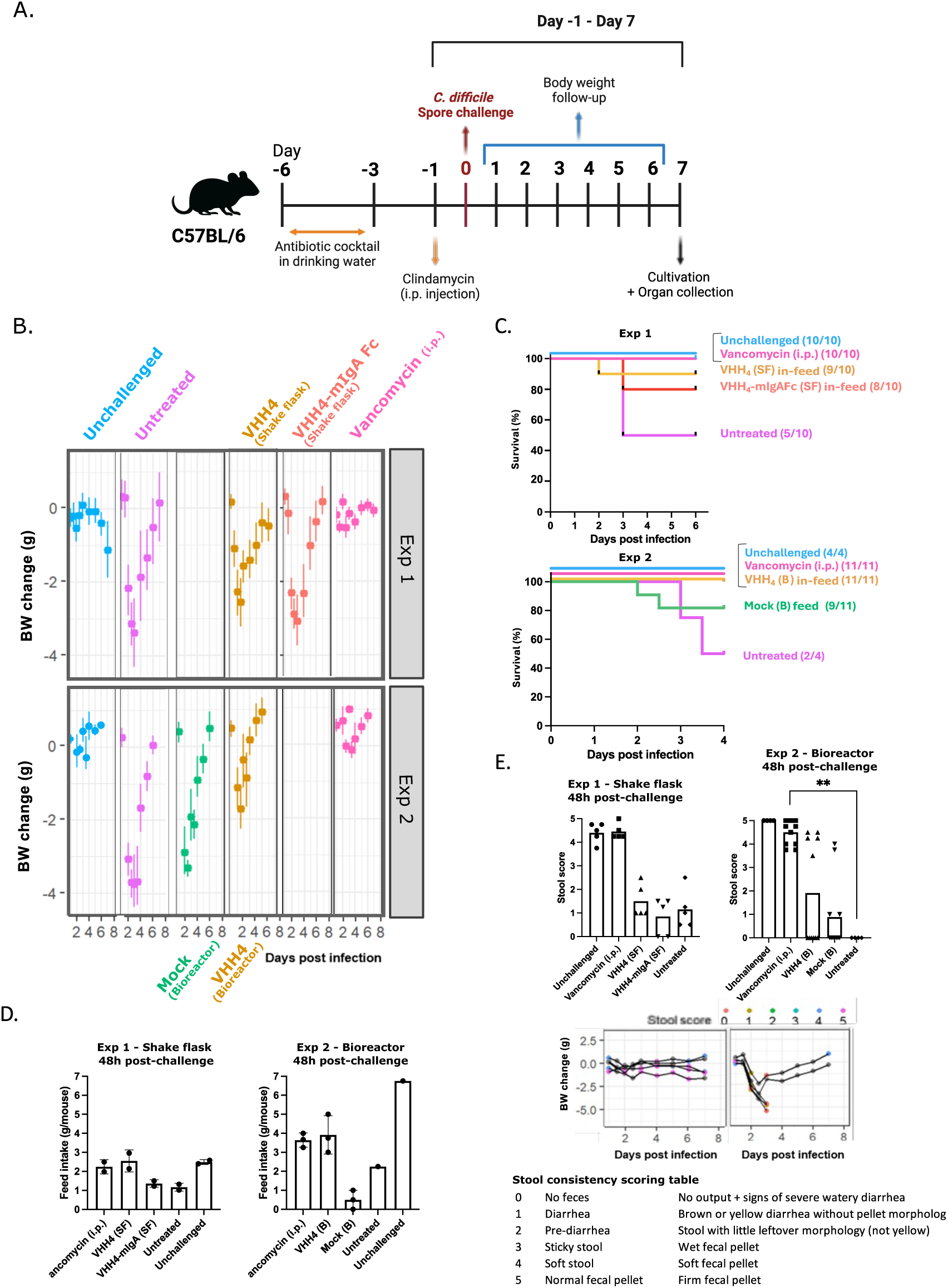
Oral administration of VHH_4_ pellets mitigates disease in a murine CDI model. (A) Experimental workflow and timeline of the *C. difficile* spore challenge model. Visual created using BioRender.com. (B) Body weight loss across groups following infection for Exp 1 (n = 10) and Exp 2 (n = 11) shows efficacy of VHH_4_ compared to Untreated or Mock (B) treated mice in both experiments. (C) Kaplan–Meier survival showing protection from mortality by treatment with VHH_4_ produced in shake flask (Exp 1) or in bioreactor (Exp 2), while VHH_4-_mIgA Fc (shake flask) (Exp 1) showed less potential. (D) Feed intake comparison across groups at 48 hours post-infection for Exp 1 (left) and Exp 2 (right) (n = 3 cages). (E) Stool scores 48 hours post-infection reveals improved clinical outcomes in vancomycin-treated mice and a non-significant improvement in VHH_4_-treated mice (Mann-Whitney – Untreated vs Treated: p = 0.0015 for vancomycin, p = ns for VHH_4_). Visual and blindly quantified stool scores correlate well with primary disease outcomes (BW change) and support efficacy findings.

*C. difficile* infected mice receiving pellets without antibodies (Untreated), infected mice receiving pellets without antibodies but treated therapeutically through daily intraperitoneal (i.p) injections (Vancomycin) and antibiotics pre-treated yet mock (PBS)-challenged healthy control mice receiving pellet without antibodies (Unchallenged) mice were taken along as controls. In our first *in vivo* mouse efficacy study, VHH_4_ (Shake flask) and VHH_4_-mIgA Fc (Shake flask) feed pellets were compared (Exp 1).

First, we found that ad libitum and continuous feeding of infected mice with pellets containing shake flask-produced VHH_4_ showed significant mitigation of the disease with reduced weight loss on days 3-4 post-challenge (VHH4 (SF) |vs| Untreated; p < 0.05) (**Fig. 3B, Exp 1**) and a lower mortality rate compared to infected mice eating pellets not containing any antibodies (90 % vs 50 % survival) (**Fig. 3C, Exp 1**). In contrast, VHH_4_-mIgA Fc did not significantly impact BW compared to untreated mice in our setup, and showed twice as much mortality (80 % survival) as compared to VHH_4_, while the standard of care therapeutic systemic antibiotic treatment-mimicking positive control (i.p. injections of vancomycin) fully prevented disease (**Fig. 3B-C, Exp 1**). These *in vivo* results indicate that the VHH_4_ pellet feed outperformed VHH_4_-mIgA Fc. Therefore, we can conclude that in this *in vivo* disease model and with the currently used formulation, the pellet-formulated VHH_4_-mIgA Fc fusion is inferior to VHH_4_ in its *in vivo* efficacy, despite much higher *in vitro* potency against TcdA of the non-feed-formulated purified antibody protein.

Feed intake was followed up during the experiment and generally the intake was lower for VHH_4_–mIgA Fc and untreated mice as compared to the other groups (**Suppl. Fig. 3G, Exp 1**). Two days post-challenge, when the infection is most fulminant, mice ingested a median of 2.55 g of VHH_4_ feed and 1.36 g of VHH_4_-mIgA Fc feed daily, which likely is due to a lack of appetite due to the developing enteric infection (**Fig. 3D** and **Suppl. Fig. 3G, Exp 1**). Finally, the average amount of feed eaten per mouse daily in the five days following challenge was calculated and again here, it is clear that VHH_4_ treatment enabled the mice to keep eating at least the same amount of food as unchallenged animals, whereas this was not at all the case for VHH_4_-mIgA Fc. (**Table 3**).

**Table 3.** Average daily feed intake and standard deviation (SD) per treatment group over the four days following *C. difficile* spore challenge (n = 2 for Exp 1 and n = 3 for Exp 2).

| Exp 1 - Shake flask (SF) |  |  |
| --- | --- | --- |
| Treatment group | Average feed intake<br>per mouse (g/day) | SD (g/day) |
| Unchallenged | 3.26 | 0.83 |
| Vancomycin (i.p.) | 2.79 | 0.56 |
| VHH<sub>4</sub> (SF) | 3.77 | 1.84 |
| VHH<sub>4</sub>-mIgA (SF) | 2.33 | 0.78 |
| Untreated | 2.41 | 1.06 |
| Exp 2 - Bioreactor (B) |  |  |
| Treatment group | Average feed intake<br>per mouse (g/day) | SD (g/day) |
| Unchallenged | 4.58 | 2.02 |
| Vancomycin (i.p.) | 3.91 | 0.62 |
| VHH<sub>4</sub> (B) | 3.74 | 1.05 |
| Mock B | 2.74 | 2.07 |
| Untreated | 3.19 | 1.06 |

Finally, fecal pellets were independently scored on a diarrheal qualitative scoring system and outcomes were found to be in line with primary disease outcomes (**Fig. 3E, middle** and **bottom**), however at 48 hours post-infection, stool scores did not significantly improve compared to untreated controls in either of the testing groups. Taken together, our findings show the first hints of efficacy of orally administered edible VHH-type antibodies in a preclinical model of *C. difficile* in mice. We suggest that - with further drug development efforts - oral delivery of unpurified, *Pichia*-produced anti-toxin VHHs could offer a non-invasive and non-antibiotic prophylactic treatment option for the (r)CDI patient population in the future.

### Scaled-Up Bioreactor-derived VHH_4_ confers robust prophylactic protection against CDI compared to a matching mock fermentate despite partial fragmentation

To scale-up production of the VHH_4_ molecule, a 30 L bioreactor (B) cultivation process was devised. A repeated fed-batch run consisting of four draw-fill cycles ran for a total of about 300 hours (12.5 days). The individual cycles can be clearly distinguished in the biomass concentration plot (dry cell weight (DCW) over time), with each drop in the curve representing a draw and fill step (**Suppl. Fig. 2C, left**). The productivity of the cells throughout the process showed an upwards trend over the cycles (**Fig. 2B** and **Suppl. Fig. 2C, middle)**. After DSP through centrifugation and ultrafiltration/diafiltration, an average yield in the harvested culture medium of 50 mg/L was obtained and, in total, almost 2 g of intact, tetrameric VHH_4_ was produced for further formulation into edible pellets for the mice (**Table 4**).

**Table 4.** Summary of VHH_4_ expression under controlled bioreactor conditions. Supernatant (SN) Volumes, Yields after Downstream processing (DSP), Amount of VHH_4_ per cycle and Cumulative VHH_4_ amount are listed. Only the intact VHH_4_ (56 kDa) band was considered for the densitometry-based protein quantification.

|  | Volume<br>SN (L) | Amount of VHH<sub>4</sub>*<br>(mg) | Estimated<br>yield* in SN<br>(mg/L) |
| --- | --- | --- | --- |
| **Cycle 1** | 10.0 | 375 | 37.5 |
| **Cycle 2** | 11.0 | 333 | 30.3 |
| **Cycle 3** | 10.3 | 541 | 52.6 |
| **Cycle 4** | 9.7 | 738 | 76.1 |
| *VHH<sub>4</sub> in its tetrameric form |  | **Cumulative amount**<br>1,987 | **Average yield**<br>49.3 |

Bioreactor-produced VHH_4_ showed significantly higher fragmentation behavior as compared to the shake flask-produced constructs (**Fig. 2D, right**). An anti-VHH immunoblot revealed 43 % of the fully tetrameric construct was left after DSP and pelleting, next to 19 % of trimeric (40-45 kDa), and 38 % of dimeric VHH (30-35 kDa) (**Fig. 2D, table**). The more pronounced fragmentation behavior in the multi-cycle bioreactor process can likely be explained by the higher shear stress in the bioreactor vs. shake flask, which results in more cell lysis and the release of proteases into the medium, leading to such clipping events.

To further fine-tune the *in vivo* efficacy experiment, we investigated whether there was an effect on mouse bodyweight evolution when they were fed pellets containing mock culture medium of *Pichia*, a condition not yet included in the first mouse efficacy study. To this effect, a palatability experiment in mice was performed using mock pellets - which contained concentrated culture medium obtained from a shake flask cultivation of the parental host NCYC 2543 *Pichia* strain that did not produce any recombinant protein. Interestingly, we observed an increased body weight (BW) gain over time in mice receiving these shake flask-produced yeast SN-based pellets (**Fig. 2E, left**). The BW increase was more pronounced in case when *Pichia* was grown on glycerol as its primary carbon source, as compared to methanol, despite the mice eating 4.5-5 g of all feeds per day on average. Notably, a transient increase in feed intake was often observed on the days following the feed tank switch (day 1-2), which is also reflected as outlier data points (**Fig. 2E, right**). As our bioreactor manufacturing process uses glycerol as carbon source, it was clear that an appropriate background-fermentate pellet feed control was warranted to fine-tune our second *in vivo* efficacy-testing experiment. To enable this production of mock pellets, wildtype *Pichia* host strain NCYC 2543 was grown in a single cycle of the fed-batch process **(Suppl. Fig. 2C, right),** which provided enough of the material. The harvested SN of the mock run was processed in the same way as for the VHH_4_-containing culture medium and concentrated by the same factor to obtain comparable host-cell protein and any remaining medium-derived macromolecular component concentrations in mock and VHH_4_-containing feed pellets.

With these bioreactor-produced feed materials in hand, we performed a second *in vivo* mouse efficacy study in which we compared the peri-exposure prophylactic efficacy of VHH_4_ (Bioreactor) produced in the 30 L bioreactor runs with a group receiving pellets containing processed SN of the mock bioreactor run (Mock (Bioreactor)), next to the other control groups also included in the earlier mouse efficacy experiment. Statistical analysis revealed here a more robust significant improvement in body weight on days 2 to 5 post-challenge compared to the mock (VHH_4_ (B) |vs| Mock (B); p < 0.05) (**Fig. 3B, Exp 2** and **Suppl. Fig. 3E**), and all mice survived infection (**Fig. 3C, Exp 2**). During this second *in vivo* study, overall feed consumption was higher for most groups than in the first experiments, as these mice had a higher average baseline BW. Remarkably and very encouragingly, the food intake of mice obtaining VHH_4_ was again comparable or better than that of Vancomycin treated mice throughout the experiment **(Fig. 3D** and **Suppl. Fig. 3G, Exp 2),** indicative of absence of enteric disease of a severity that would make mice eat less. Double blind stool scoring by two volunteers in Exp 2 detected almost half of the population (5/11 mice) receiving VHH_4_ showed improved stool scores 48-hours post-infection resulting in a slight but non-significant improvement of this secondary outcome (Mann-Whitney – Untreated vs Treated: p = 0.0015 for vancomycin, p = ns for VHH_4_) (**Fig. 3E, top**). Considering both experiments, peri-exposure prophylactic administration of VHH_4_ did significantly reduce mortality compared to untreated mice (HR 0.09, p = 0.027) (**Suppl. Fig.3F**).

## Discussion

In this study, we have demonstrated that oral administration of a recombinant, non-purified, *Pichia*-produced tetravalent VHH-type antibody (VHH_4_) can improve disease outcomes in a preclinical murine model of *Clostridioides difficile* infection (CDI). While the VHH_4_ used throughout this manuscript has already shown *in vivo* potency by Chen *et al.* when *in situ* expressed in the form of an IgG fusion molecule by *Saccharomyces boulardii*^29^, here we have shown that the non-fused tetrameric VHH also improves disease outcomes when administered after being dried into murine feed pellets. A live cell approach shows good efficacies and seems revolutionary – engineering commensals for cost-effective *in situ* production of biotherapeutics - yet some limitations include long-term storage and safety, as exact doses of exogenous antibody secretion will be hard to estimate or control^35,36^. Moreover, this strategy presents regulatory challenges in some parts of the world, as it involves the administration of a Genetically Modified Organism (GMO) to patients. The fact that the protein can be formulated in a dried, stable, safe and easily administered form with more controlled dosing capabilities as compared to *in situ* expression, and still provide evidence of efficacy, is one of the main outcomes of this study.

Even though *in vitro* results seemed in favor of VHH_4_-mIgA Fc as the more potent molecule (1.316 U/nmol against the TcdA/B mix) as compared to the VHH_4_ molecule (0.032 U/nmol against TcdA/B mix), this did not translate into promising *in vivo* outcomes. Therefore, it is clear that the gained *in vitro* potency against TcdA by introducing bivalency due to the mIgA Fc fusion, as seen with purified protein in the cellular assays, does not translate to enhanced feed pellet-formulated efficacy in our murine CDI model. Whether the more complex VHH-Fc fusion protein fold suffers more from the formulation and drying process compared to the small and stable VHHs, resulting in greater loss of toxin-neutralization potencies, or whether a physiological reason lies behind this observation, remains to be investigated.

In the explored tetrameric VHH, the VHH modules are linked by flexible (G_4_S)_3_ linkers, which can be prone to protease attack. Indeed an increase in protein clipping occurred upon higher cell density fermentation, likely due to more shear stress, resulting in partial cell lysis and concomitant release of proteases into the medium^39^. This fragmentation behavior, which could be directed to the linker regions, or against other labile regions in the VHHs such as the variable complementarity-determining regions (CDR) loops, has led us to investigate the activity of the individually recombinantly engineered monomers, more thoroughly. Interestingly, we have confirmed that the recombinant monomers are also capable of neutralizing either TcdA or TcdB, yet at a 50 - 5,000 -fold lower potency^28^. Therefore, we theoretically assumed that the potency of the intact molecule will be far more superior and that clipping products in the mixture will not have a huge impact on *in vivo* outcomes. Interestingly, bioreactor-produced VHH_4_ even exhibited a more robust prophylactic effect in our murine CDI model, a subject for further study.

We decided to formulate the feed pellets at 10 nmol antibody per gram of feed in the present study, assuming mice eat about 3-4 g per day and would ingest 30-40 nmol or 1 U, enough to theoretically neutralize 50 % of the cytotoxicity caused by 1 nmol TcdA or TcdB in our *in vitro* assays. This equals a current dose of ± 85 mg/kg for mice. When calculating the human equivalent dose (HED), this results in 7 mg/kg in humans, or a daily dose of about 500 mg of antibody for a 70 kg person^40^. A follow-up experiment should include a dose-escalation study in mice, or in more translational models for GI infectious diseases such as gnotobiotic pigs^41^. In this study, average titers of approximately 50-100 mg/L were obtained from a non-engineered wildtype *Pichia* strain using a standard production process. We are confident that, using common techniques used in *Pichia* to increase the yields, including optimal promoter and secretion signal choice, copy number increase, chaperone overexpression, use of protease knockout strains, and thorough process optimization, a cost-effective and scalable production platform for this oral antibody prophylactic therapeutic lies within current possibilities^42–44^.

In our work, the fermentation process was streamlined with an at-scale down-stream processing protocol including uf/df and buffer exchange steps, and a subsequent formulation method using pellet extrusion, which is straightforward to implement in any standard biopharmaceutical or food/feed production unit. Dosing of the antibodies into the feed pellets for mice experiments allows for accurate dose determinations as feed intake is easily monitored by daily weighing of the feed tank. One limitation to our in-feed administration method is that mice that were more affected by the *C. difficile* spore challenge, also ate less, inevitably leading to a loss of intake of the VHH_4_ toxin-neutralizing molecule. This potentially has increased the difference in outcome between both the VHH_4_ and VHH_4_-mIgA Fc format. For this preclinical evaluation in mice, we have extruded and dried the antibodies for 20 hours at 40 °C. In previous work on ETEC we have used freeze-drying of the antibody-containing feed material, a method that is generally much friendlier to protein integrity, and an obvious alteration for future work upon translation towards clinical experiments.

The outcomes using VHH_4_ derived from either shake flask or bioreactor productions, gave similar positive *in vivo* outcomes in our experiments. This indicates that further bioreactor scale-up is technically feasible. Such therapy should ideally be used preventively, for example handed to an at-risk individual while broad-spectrum antibiotics is given for general surgery, or in larger scale during hard-to-control hospital outbreaks of *C. difficile*. Additionally, no chromatography is needed, making the full production process more economical. At a larger scale, and for human therapy, a DSP including spray-drying and fluid-bed steps for micro pelleting and simultaneous enteric coating, such yeast supernatant-containing powders or capsules could be ideal, not only for (r)CDI, but also for emergency situations and outbreaks of enteric pathogens that can be neutralized by VHHs.

Taken together, our data suggest that with further drug development efforts, especially in finding the minimal effective dose (MED) and engineering of both the strain and production process for optimal recombinant protein production of lead molecule and its in-feed formulation, oral delivery of *Pichia*-produced anti-toxin VHHs could offer a non-invasive and non-antibiotic preventive treatment option for the (r)CDI patient population.

## Materials and Methods

### Construction, expression and purification of VHHs and VHH-Fc’s in *Pichia*

Wildtype *Komagataella phaffii* (NCYC 2543) was chosen as an expression host for this study. The TcdA/B-neutralizing antibody lead, VHH_4_, was selected from literature^45^. The VHH_4_-mIgA Fc fusion antibodies were constructed by grafting the VHH_4_ coding sequences N-terminally to the hinge region of the native sequence of murine IgA. The genes of interest were in frame with an N-terminal secretion leader (Ost1 or Kar2) and C-terminally fused to a HIS_8_ tag, which allowed straightforward downstream affinity purification and analysis. The expression plasmids were constructed with an in-house modular cloning protocol for yeast (MoClo), based on Golden Gate cloning^44^. After transformation in *Pichia,* clone screening and small-scale expression tests, the antibodies were expressed in 2 L baffled shake flasks for 48 h in BMGY, under standard conditions, and purified. For purifications, the filtered supernatant was loaded onto a HisTrap Excel (5mL) column (Cytiva). After washing (5 column volumes (CV) of 20 mM imidazole, 500 mM NaCl, 20 mM NaH2PO4/Na2HPO4, pH 7.4), the bound proteins were eluted (10 CV, 400 mM imidazole, 500 mM NaCl, 20 mM NaH2PO4/Na2HPO4, pH 7.4) and analyzed on SDS-PAGE. Afterwards, the fractions containing the protein of interest were pooled and loaded on a HiLoad 16/600 Superdex 200 pg column (Cytiva) equilibrated with PBS buffer (Phosphate Buffered Saline, pH 7.4). Eluted fractions were analyzed on SDS-PAGE. Selected fractions were pooled and finally concentrated in PBS buffer over a 3 kDa cut-off centrifugal filter unit (Merck Millipore, UFC901008). The protein concentration was calculated by measuring the absorbance at 280 nm.

### Large-scale shake flask expressions

For large-scale shake flask expressions, a 5 mL preculture of the expressing clones was inoculated in BMGY medium supplemented with 100 µg/mL Zeocine. After 24h of growth and after checking the inoculum for bacterial contamination under a Light Microscope, 20x 2 L baffled shake flask containing 250 mL BMGY were inoculated with the preculture aiming for a start OD of 0.1. Production was allowed for 48 hours at 28°C, on a shaking platform set to 200 RPM. The supernatant was harvested by centrifugation (10,000 G, 15 min, 4 °C), filtered over a .22 µm bottle top filter, and finally pooled.

### Bioreactor cultivation and VHH_4_ expression

All fed-batch cultivations were carried out in a Biostat® Cplus (Sartorius) 30 L bioreactor. The process commenced with the batch phase at 10 L of medium containing 0.5 g of MgSO_4_·7H_2_O, 1.83 g of citric acid, 12.4 g of (NH_4_)_2_HPO_4_, 0.9 g of KCl, 0.022 g of CaC_l_2·2H_2_O, 40 g of glycerol, and 10 g of yeast extract per liter distilled water, with the pH adjusted to 5.5 using HCl. The medium was sterilized in place, and once cooled, 4.6 mL/L of temperature-sensitive PTM1 trace salt solution was aseptically added. The process was carried out at 25 °C and at a pH of 5.0 (adjusted with the addition of 30% ammonium hydroxide solution (NH₄aq)), employing 1 vessel volume per minute (vvm), and maintaining a 30% dissolved oxygen (DO) level throughout. A customized dissolved oxygen (DO) cascade was implemented to keep the DO level at the set point, beginning with an increase in agitation from 600 revolutions per minute (rpm) to 1200 rpm. Once agitation reached its maximum limit, the sparged gas stream (which remained steady at 1 vvm) was supplemented with pure oxygen to the necessary level, typically required towards the end of the Batch phase and during the Fed-Batch Phase.

The Batch Phase was initiated with the addition of the inoculum with a theoretical starting optical density at 600 nm (OD600) of 0.8. *Pichia* expression strains generated for VHH_4_ expression were evaluated in repeated fed-batch cultivation mode (4 cycles). For background NCYC2543 strain cultivation, which serves as ock, a classical fed batch fermentation was run (1 cycle). For the inoculum, the strains were grown in a baffled 1 L shake flask containing 100 mL of BMGY medium at 28°C with 200 rpm agitation until an OD600 of 20-40 was reached. Before inoculating the bioreactor, the inoculums were inspected microscopically to confirm the absence of contamination. The Fed-Batch Phase began following the detection of the DO spike by introducing 8.0 L of Feed Solution (50 % of glycerol v/v) with a linearly increasing feeding profile over about 70 hours. The glycerol concentration in the culture broth remained at limited levels throughout this phase, which was verified by the metabolite analysis of the samples via HPLC. This feeding strategy resulted in a gradual decrease in specific growth rates over time. With the addition of 8.0 L of Feed Solution and about 0.750 L of Base solution to the 10 L of batch medium, the volume reached the maximum operational capacity of the vessels, 16 L of cell broth was removed from the vessel to serve as partial harvested material, and 8.0 L fresh medium was added to the 2.0 L remaining live cell broth, serving as the preculture for the next cycle, which followed the exact same procedure as just described for the first cycle. The cells were harvested through centrifugation (10.000 G for 15 min), followed by clarifying the supernatant using bottle-top vacuum filtration (0.22µm PES). Harvests from four cycles were kept separately for intermediate evaluation of the proteins during these first DSP steps.

Mock runs using the wildtype NCYC 2543 background strain were performed in the same manner as described above for VHH_4_ runs, however with the following adaptations: A single cycle fed batch fermentation was performed with a starting OD in the batch phase of 0.02 OD600 values, after which a linearly increasing feeding strategy was applied in the next 52 hours FB run time. The full broth of 17 L was collected at the end of the fed-batch phase, after which the vessels were cleaned using standard procedures. Samples from different phases of the process were preserved at – 20 °C for subsequent analysis.

### Viability check and sample processing during the bioreactor cultivations

Broth samples were collected from the bioreactor at regular intervals during the batch and fed-batch phases. Samples were analyzed for biomass concentration via optical density (OD) and Dry Cell Weight (DCW) measurements in triplicates. For DCW measurements, a moisture analyzer (MA35, Sartorius) was used. All fractions were frozen in liquid nitrogen and stored at -20 °C between every analysis.

### Characterization of *Pichia*-produced antibodies by SDS PAGE

To characterize the recombinantly produced proteins, reducing or non-reducing SDS-PAGEs were run. In some cases, an endoglycosidase H (endoH-HIS_8_, made recombinantly in our lab) digest was performed to trim down N-linked glycans, prior to analysis. For that purpose, the pH was adjusted using a 0.1 M acetate buffer, pH 5.5 and the samples were boiled in a heat block at 98 °C for 10 min. 1 µg of endoH was added after cooling on ice and incubation at 37 °C for 10 hours. For reducing SDS-PAGE, 5 µL of 5x Laemmli buffer with dithiothreitol was added to 20 µL of sample and the samples were boiled in a heat block at 98°C for 10 min. 15-25 µL was loaded on 4-20% gradient polyacrylamide gels (GenScript ExpressPlus, M42015). The gels were either stained with Coomassie Brilliant Blue R250 for at least 1 hour before destaining with a 20 % (v/v) methanol and 10 % (v/v) acetic acid in water solution, or blotted on a nitrocellulose membrane (1 hour, 150 V). For detection of the HIS_8_ tag, a 1:10,000 dilution of DyLight800-coupled anti-HIS_8_ tag antibody (Rockland, 33D10.D2.G8) was used. For detection of the VHH domain, a 1:10,000 dilution of a biotinylated AffiniPure® Goat Anti-Alpaca IgG (Jackson Immuno Research Labs, 128-065-232) was used followed by incubation with a DyLight800-coupled Streptavidin Protein (Thermo Scientific, 21851). Fluorescence was recorded at both 700 nm and 800 nm.

### Characterization of *Pichia*-produced antibodies by ELISA

The presence of the anti-TcdA/B constructs in the different biological samples was assessed by HIS-capture ELISA. Wells of a microtiter plate (MaxisorpTM 96-well plates, Nunc, 442404-21) were coated overnight at 4 °C with Monorab Rabbit Anti-Camelid VHH antibody (Genscript, A01860-200) at a concentration of 200 ng/well, diluted in 50 mM NaHCO3/Na2CO3, pH 9.6. After washing four times with PBST (0.05% (v/v) Tween20 in PBS), 200 μL of blocking buffer (3 % milk powder in PBST) was added and the microtiter plate was incubated for two hours at room temperature. Intestinal samples were thawed, centrifuged (10,000 G, 10 min, 4 °C) and diluted 1 in 5 in sample diluent (0.05% Tw20 (v/v), 1x protease inhibitor (complete P.I., Roche) in PBS) before adding 100 µl of samples to the wells. In addition to a standard curve of purified protein at known concentrations ranging from 200-0.001 nM (1:3 dilutions), four spike controls of purified protein in the corresponding matrix background (0.741 nM, 0.247 nM, 0.082 nM, 0.027 nM) were also included in order to evaluate the matrix effect. Incubation was continued overnight at 4 °C. The wells were washed four times with PBST and then incubated with a goat anti-alpaca IgG-HRP-conjugated antibody (Jackson Immuno Research Labs, 128035232, 1/25,000 dilution). After one hour of incubation and washing with PBST, 3,3′,5,5′-tetramethylbenzidine (TMB) was added as substrate and absorbance of the solution in the wells was measured at 450 nm with an ELISA reader after adding H_2_SO_4_ (2N) to stop the reaction.

### Vero cell toxin-neutralization assay

Vero cells were purchased from ATCC (No. CCL-81). The day before the start of the experiment, cells were counted and seeded at 5E3 cells per well in 96-well culture plates. Three-fold dilutions of neutralizing VHH-containing constructs were pre-incubated with a specific amount of toxin determined during optimizations assay (15-30-fold the IC50): 50-100 ng/mL for TcdA, 0.05-0.1 ng/mL for TcdB, and a 1:1 equipotent mix for TcdA+B. Native toxins were purchased from The Native Antigen Company (CDA-TNL-50, CDB-TNL-50) as a lyophilized powder. Mixtures were incubated for 1 hour before being added to the cells which had been growing for 24 hours. The plates were further incubated at standard growth conditions over the next 48 hours. Using a light microscope, cells were visualized at different timepoints after challenge (0h, 3h, 16h, 24h, 48h) in order to observe cell-rounding events. Two days after toxin challenge, wells were washed three times with PBS and the viable cells were quantified using ATP-based luminescence reporter assay as a readout (CellTiter Glo® Lumiscent Cell Viability Assay, Promega). Final assays were repeated in triplicate and average IC50 values were determined by non-linear regression on the sigmoidal curves (Graphpad Prism).

### PK experiments

Healthy male C57BL/6 mice (Charles River Labs) between 8-10 weeks old were used for this experiment. The animals were housed in a temperature-controlled environment with 12 h light/dark cycles; food and water were provided *ad libitum*. All experiments were done under conditions specified by law and authorized by the Institutional Ethical Committee on Experimental Animals (Ethical application EC2019-033). The study consisted of 5 experimental groups with 6 mice per group. Two groups received 100 µL of anti-TcdA/anti-TcdB VHHs and 2 groups received anti-TcdA/anti-TcdB VHH-IgAs *per os (p.o.)* at a dose of either 1 mg/mL or 0.1 mg/mL. The control group received the same amount of a mock vehicle formulated in the same manner yet not containing any experimental protein. Prior to administration of these test compounds, mice received 100 µL of a gastroprotective vehicle *per os* (p.o.) by gavage, consisting of 40 mg/mL milk solution in 0.1 M NaHCO_3_. Three hours post-dosing, three animals were sacrificed by injection of a mixture of ketamine (10 %) and xylazine (2 %), and intestinal samples were collected according to the anatomical borders (luminal content and homogenates of stomach, small intestine, caecum, colon). Additionally, feces and blood samples were collected by the clean catch method and via jugular vein bleeding, respectively. Feces were collected in pre-weighed tubes and resuspended 20x (w/v) in PBS containing protease inhibitors (cOmplete P.I., Roche). Serum was prepared from whole blood after addition of protease inhibitors (Sigmafast) and clotting for 30 minutes at RT, followed by centrifugation at 3,500 RPM for 15 min at 21 °C and collection of the supernatant fraction. At 7 hours post-dosing, the final three animals per experimental group were sacrificed and sampled in the same manner as described above.

### Downstream Processing (DSP)

At the end of the fermentation process,10-20 L of harvest broth was clarified by centrifugation (10.000 G, 15 min., 4 °C) and filtered using the Sartojet^®^ pump with a .22 µm filtration cassette. The filtrates were further concentrated to ± 1.0 L, and consecutively 6x buffer exchanged with a 17.5 mM NaCl containing sodium phosphate buffer (20 mM NaPi Low Salt, pH 6.0). A 6x buffer-exchange should theoretically result in a > 99 % complete replacement of soluble buffer components < 5,000 g/mol surrounding the proteins of interest. Finally, the buffer-exchanged medium was further concentrated through a semi-automated crossflow filtration system (MWCO 10 kDa) to a given volume based on the desired final concentration per gram feed pellet (SartoFlow®, Sartorius).

### Formulation and gamma irradiation of unpurified SN pellets

The concentrated protein mixture (for experimental groups) or low-salt buffer (for control groups) is homogenously admixed to powdered mouse feed (SNIFF) using a Kenwood Chef Food Processor in a pre-determined ratio of 700 mL/kg feed and hereafter extruded and cut into pellets (2.5 cm x 1.0 cm). Mouse feed pellets were air dried for 20 hours at 40 °C in a food dehydrator (Klarstein). Next, batches of pellets are γ-irradiated with 5 kGy spread over 5 hours to reduce heat and ray intensities and avoid breakdown or fragmentation of the POI as has been described^46^. Irradiations were performed at the BR2 Co60 Brigitte underwater irradiation facility of the SCK CEN (Mol, Belgium). Pellets were stored in airtight Ziplock bags in the dark at room temperature for a maximum of one month before the start of the experiment.

### Spore cultivations

*Clostridioides difficile* strains were bought from ATCC. Both VPI-10463 (Ribotype 078) and Strain 630 (Ribotype 012) were tested for sporulation efficiency. The protocol used for *C. difficile* spore isolation is largely based on work by A. N. Edward *et al.* (2016)^47^. In short, in an anaerobic chamber containing a mixture of 5 % CO_2_, 10 % H_2_, 85% N_2_, vegetative *C. difficile* was grown overnight in 10 mL of pre-reduced BHIS+TA+Fructose. The next day, when the culture was in active growth (OD < 1.0), 250 µl of the preculture was spread on fresh, pre-reduced 70:30 agar plates (63 % Peptone (BD Bacto), 3.5 % Proteose peptone (BD Bacto), 1.06 % Tris base, 0.7 % Ammonium sulfate, 11.1% Brain Heart Infusion (BHI) (BD Bactro), 1.5 % Yeast extract (BD Bacto), 16 % Agar (BD Bacto), 10 % L-cysteine (Sigma, C7352)) and incubated anaerobically at 37 °C for 24-48 hours. After sporulation, three randomly picked colonies were validated to be *Clostridioides difficile* by MALDI-MS before continuing the protocol. Plates were removed from the anaerobic flow and the bacterial lawn scraped off and collected in a falcon tube containing 10 mL 1x PBS. The cells were centrifuged (8,000 G, 10 min, RT) and washed once with 95% ethanol and two more times with 1x PBS. Finally, the pellet was resuspended in 10 mL 1x PBS + 1% BSA and aliquoted in glass vials. Final spore stocks were heated to 70 °C for 20 minutes and cooled down to RT for further storage until use. Whenever spores were needed for either quality control (spore counts) or *in vivo* experiments, the stock was re-heated to 70 °C for 20 min to reduce the presence of live cells in the stock.

### *In vivo* proof of concept experiments

11-12-week-old female C57BL/6 were used for these experiments (Charles River France). Animals were housed with 3-5 mice per cage in a temperature-controlled environment 12 h light/dark cycles; food and water were provided *ad libitum*. All experiments were done under conditions specified by law and authorized by the Institutional Ethical Committee on Experimental Animals (Ethical applications: EC2019-033, EC2024-055). All experiments were performed in a Biosafety Level 2 facility of the animal house facility. All mice received an antibiotic cocktail in their drinking water for 3 days to induce microbiota dysbiosis (Kanamycin (0.4 mg/mL), Gentamycin (0.035 mg/mL), Colistin (850 U/mL), Metronidazole (0.215 mg/mL), Vancomycin (0.045 mg/mL)). Thereafter, mice were put on normal drinking water for 48 hours. The day before the challenge, an i.p. injection of the broad-spectrum antibiotic Clindamycin (10 mg/kg) was given to the mice for a complete eradication of the endogenous microbiota. During this preparative phase, stool samples were collected to verify full eradication. Therefore, fecal pellets were homogenized in 1 mL sterile PBS, centrifuged, and 100 µl grown in BHIS medium for 24 hours in a shaking incubator at 37 °C. Finally, OD_600_ was measured to verify dysbiosis in antibiotic-treated mice. One day after the end of the antibiotic treatment, animals were intragastrically challenged with 1x10^4^ spores of strain VPI-10463 (RT078) unless otherwise mentioned. The unchallenged, healthy control group received oral gavage of PBS instead of spores. The comparator group received daily i.p. injection of 100 µl clinically approved Vancomycin (10 mg/mL) antibiotics, with the first injection taking place at least one hour before spore challenge. Both comparator and control groups were fed *ad libitum* with a mock feed (no antibodies present in feed), switched 24 hours before spore challenge and given throughout the experiment. Feed intake was followed up once or twice daily in the following 5-7 days and the mice were scored on body weight, mortality, stool consistency and overall behavior. On the harvest day, all experimental mice were sacrificed by cervical dislocation and terminal blood was taken. Fecal samples were taken on day 0, 1, 2, 3, 4 and 5 and were scored double blinded in duplicate in Exp 2 on days 2 and 3 post-challenge.

### Statistics

The body weight data were analyzed per experiment to enable the comparison of different treatment groups occurring in each experiment. We first selected the data pertaining to days 0 to 6 (last day of follow up in Exp 2) or 7 (last day of follow up in Exp 1). We used R to model the body weight at days 1-6 or 1-7 with a mixed model using a random intercept for each individual mouse (we did not have enough observations to fit a model with a random slope). As fixed effects, we used the body weight at day 0, the treatment and for the day, a natural spline with 4 degrees of freedom (3 internal knots), in order to get a flexible fit for the time. We also allowed for an interaction between Day and Treatment so that the shape of the time-course could differ between treatments. All of this was done in R using the nlme package (to fit the models) together with the splines package (to define the natural spline for the Day variable). The quality of the fit was assessed via residuals analysis of the transformed residuals (residuals and predictors were transformed with the inverse of the Cholesky decomposed covariance matrix^48^). The significance of the Treatment variable was tested with a likelihood ratio test of the full model versus the model without the Treatment variable. Contrasts and their confidence intervals were constructed using the contrast package with the confidence interval type set to ‘simultaneous’ and the covariance type set to heteroscedasticity consistent (vcovHC). For the survival analysis, we analyzed the data from experiment Exp 1 and Exp 2 together using R. The maximal follow-up time was 7 days in Exp 1 and 6 days in Exp 2. An event was defined as obligatory euthanasia due to a body weight loss of 20% or more relative to the body weight at the start of the experiment. A mouse was considered to be censored when it was not euthanized by day 6 (Exp 2) or day 7 (Exp 1). For this survival analysis, we only considered the untreated mice and the 4x VHH or vancomycin treatment groups. These were all the groups that were present in both experiments. A stratified Kaplan-Meier plot (by experiment) was made with the ggsurvfit package and a (stratified) log rank test was performed using the survival package. Next, we performed a stratified Cox proportional hazards model with the treatment group as the predictor and the experiment (Exp 1 or Exp 2) as the stratification variable. ‘Untreated’ was the reference level so the results must be interpreted as the hazard ratio of a particular treatment group versus untreated. Note that with no events in the vancomycin group, the hazard ratio and corresponding p-value cannot be calculated. This Cox model was also run with the survival package and the results were presented with the gtsummary package.

## Acknowledgments

We would like to thank the Protein Service Facility (PSF) of VIB for their technical assistance and providing access to fermentation and DSP equipment needed for this study, Prof. Filip Van Immerseel and Dr. Venessa Eeckhaut for the use and technical support of the anaerobic chamber used for *C. difficile* spore cultivations, and Prof. Simone Beccattini for kindly providing essential protocols and strains needed for the set-up of these models in our lab.

This research was funded by a Strategic Basic Research fellowship granted by the Belgian government Fund for Scientific Research Flanders (FWO), an Industrial research fund (IOF) Advanced grant provided by Ghent University and a Strategic Valorization Proof of Concept (PoC) Project granted by VIB. The research was in part funded by VIB and Ghent University institutional resources.

