## Supplementary Figures for "ENGINEERING AND PRECLINICAL EVALUATION OF DRIED FOOD-FORMULATED YEAST-SECRETED ANTIBODIES AS PERI-EXPOSURE PROPHYLAXIS AGAINST CLOSTRIDIOIDES DIFFICILE DISEASE"

Roels *et al.*

### Supplementary material

**Supplementary figures**

A.

| Protein | Specificity | Plasmid | Expression strain | Molecular weight (kDa) | Extinction coefficient | Yield after purification (mg/L) |
|---|---|---|---|---|---|---|
| **4xVHH** | TcdA/B | PGAP, Kar2, His8, Zeo | NCYC 2543 | 56.53 | 81.415 | 6.0 |
| **4xVHH-mIgAFc** | TcdA/B | PGAP, Kar2, His8, Zeo | NCYC 2543 | 165.92 | 111.490 | 0.2 |
| **4xVHH** | TcdA/B | PUPP, Kar2, His8, Zeo | NCYC 2543 | 56.53 | 81.415 | 89.7 |
| **4xVHH-mIgAFc** | TcdA/B | PUPP, Kar2, His8, Zeo | NCYC 2543 | 165.92 | 111.490 | 53.2 |

B.

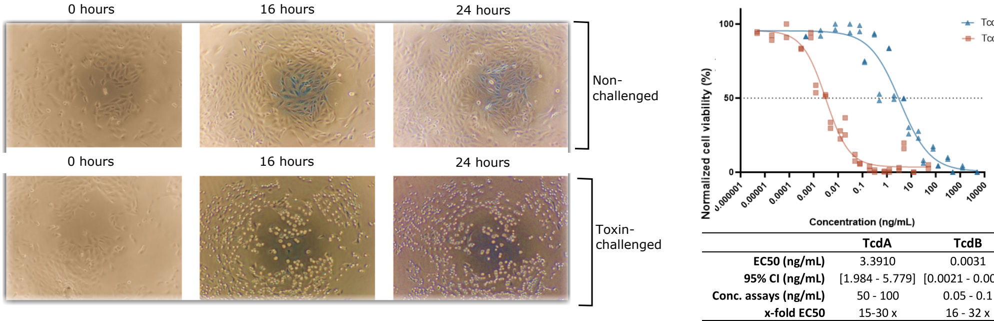


| | TcdA | TcdB |
|---|---|---|
| **EC50 (ng/mL)** | 3.3910 | 0.0031 |
| **95% CI (ng/mL)** | [1.984 - 5.779] | [0.0021 - 0.0041] |
| **Conc. assays (ng/mL)** | 50 - 100 | 0.05 - 0.1 |
| **x-fold EC50** | 15-30 x | 16 - 32 x |

C.

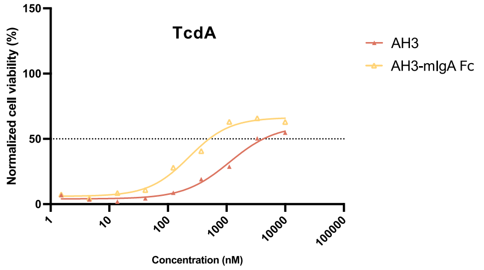


| | Toxin specificity | IC50 (nM) (Mean) | Potency mU/nmol |
|---|---|---|---|
| **AH3** | TcdA | 1,130 | 0.07 |
| **AH3-mIgA Fc** | TcdA | 230 | 0.35 |

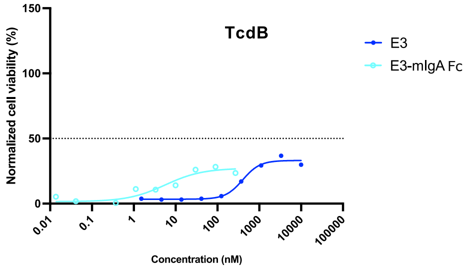


| | Toxin specificity | IC50 (nM) (Mean) | Potency mU/nmol |
|---|---|---|---|
| **E3** | TcdB | 402 | 0.00 |
| **E3-mIgA Fc** | TcdB | 6 | 0.07 |

**Supplementary Figure 1. Specifications for production and set-up of toxin-induced cell death assay for potency testing of monomers**

**(A)** Titers after uncontrolled SF *Pichia* expression improved upon switching from PGAP to PUPP and ranged from 6-90 mg/L for VHHs and 0.2-53 mg/L for mIgA fusions.
**(B)** Time-dependent toxin-induced cell-death could be visualized by increasing cell rounding events under a light microscope (Left). The concentration causing 50% cell death (EC50) was 1000-fold lower for TcdB as compared to TcdA (right).
**(C)** Vero cell-based luminescence assay showing neutralization potency of monomers against TcdA (left) and TcdB (right) showed significant lower *in vitro* activities compared to terameric constructs (n = 1 biological replicate).

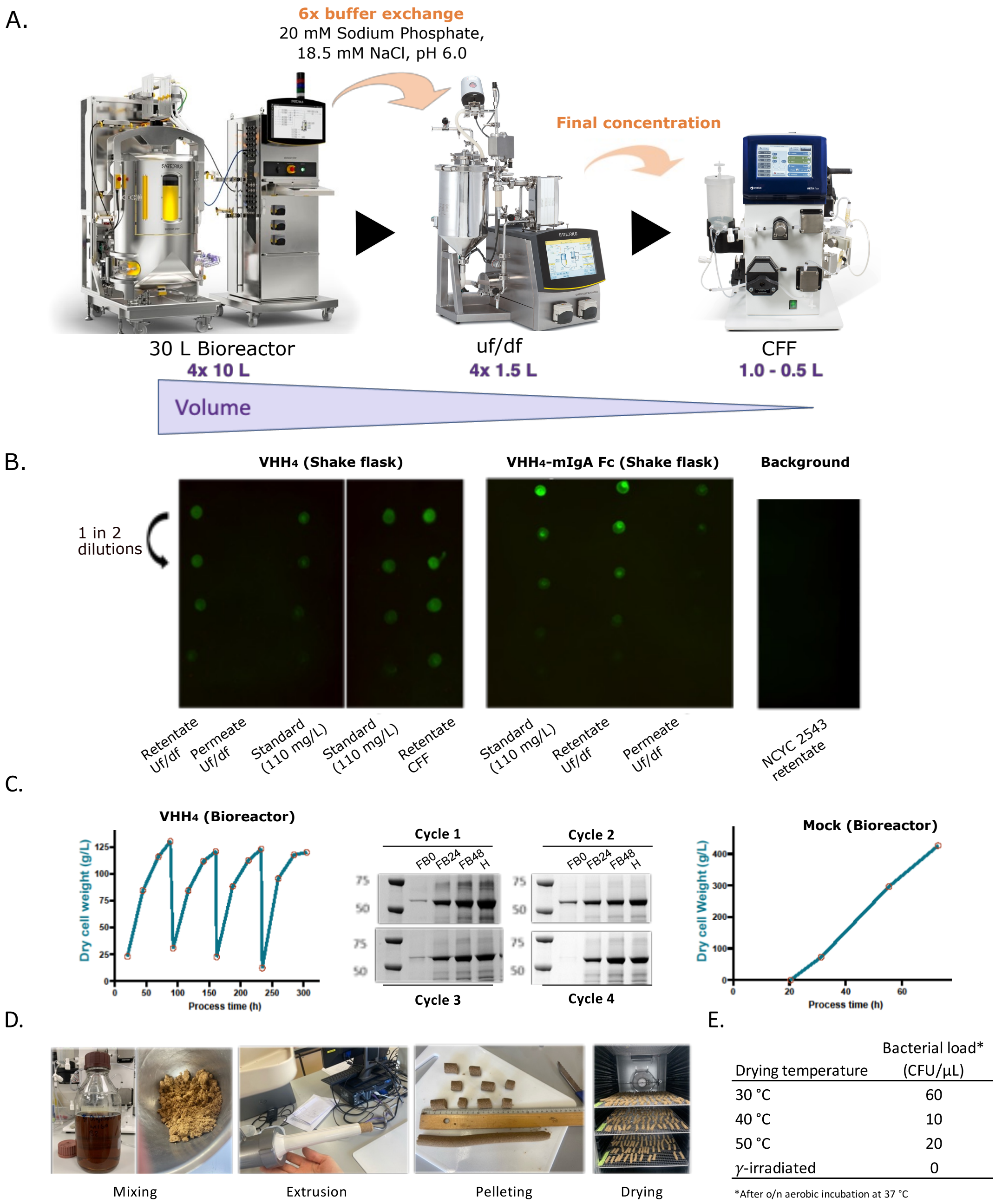


| Drying temperature | Bacterial load* (CFU/µL) |
|---|---|
| 30 °C | 60 |
| 40 °C | 10 |
| 50 °C | 20 |
| $\gamma$-irradiated | 0 |

*After o/n aerobic incubation at 37 °C

**Supplementary Figure 2. Production, downstream processing and formulation optimization of *Pichia*-produced VHH$_4$ for oral administration.**
**(A)** Overview of the downstream processing pipeline: production of VHH$_4$ in bioreactors followed by ultrafiltration/diafiltration (uf/df) and crossflow filtration (CFF) to concentrate and buffer exchange protein-containing supernatant. Images adapted from Sartorius website (*www.sartorius.com*).
**(B)** Dot blot analysis using anti-HIS detection showing concentration of HIS-tagged VHH$_4$ during uf/df and minimal loss to permeate across DSP steps.

**(C)** For the 300h VHH4 (Bioreactor) production run, four fed-batch cycles can clearly be distinguished by the biomass profile monitored by dry cell weight (DCW), indicating consistent yeast viability (left), whereas SDS PAGE confirmed consistent productivity of VHH4 in all four cycles (middle). To serve as mock vehicle, wildtype NCYC 2543 was grown in a standard (single cycle) fed-batch process (right).

**(D)** Pellet formulation process consists of mixing, extrusion, pelleting, and air drying, followed by γ-irradiation for sterilization.

**(E)** Bacterial load determined by CFU count after drying at various temperatures or γ-irradiation, showing full sterilization with irradiation and reduced microbial load at higher drying temperatures. Plates aerobically cultivated for 48 hours at 37 °C on YPD.

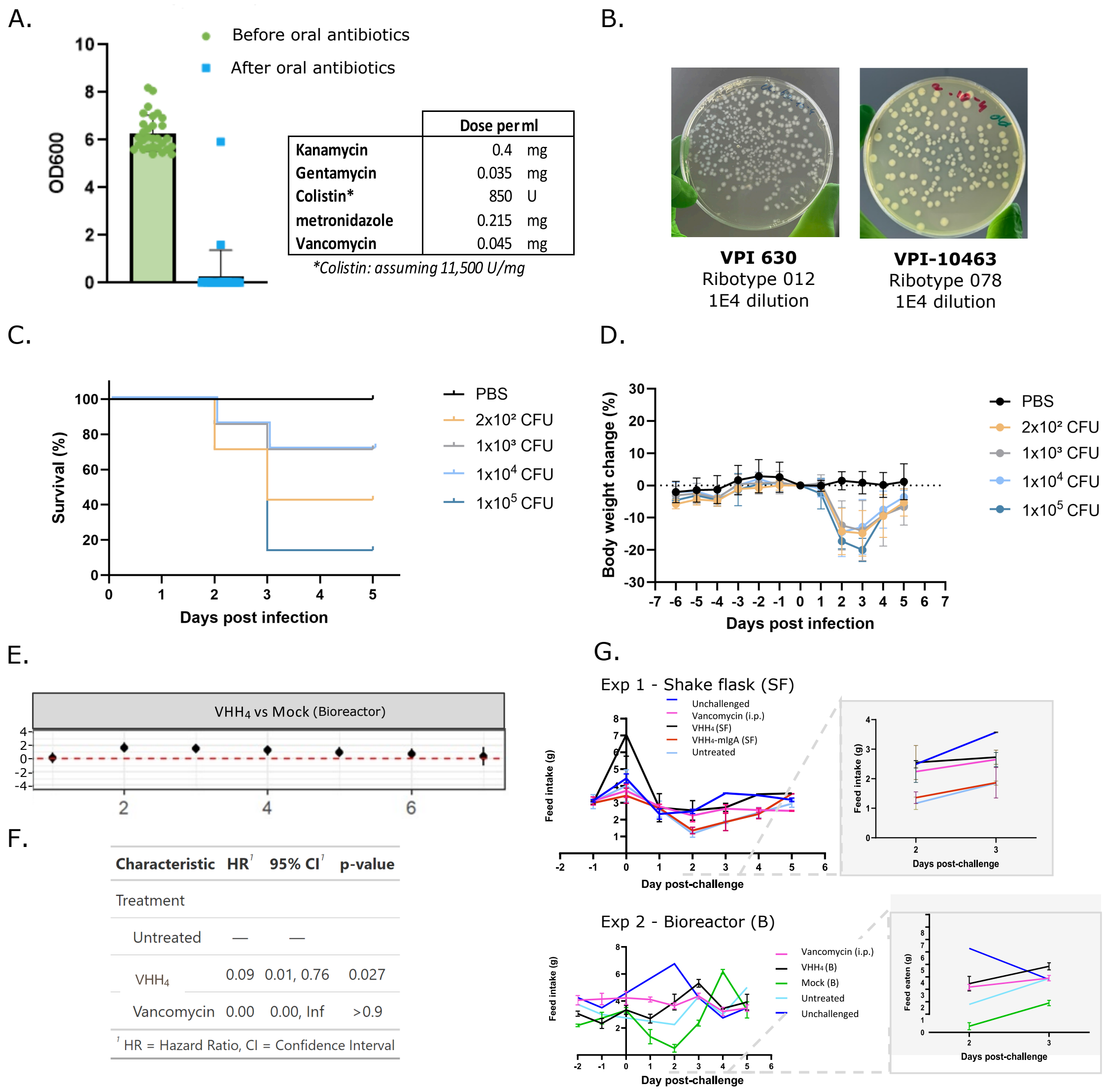


**Supplementary Figure 3. Supporting information and statistical analysis of the murine *C. difficile* challenge studies.**

**(A)** Antibiotic pre-treatment in drinking water efficiently depletes the murine gut microbiota, as shown by significantly reduced OD600 values of cultured fecal samples after 24 h low-oxygen incubation at 37 °C in Brain heart infusion-supplemented (BHIS) medium.
**(B)** Comparison of sporulation efficiencies of two clinically relevant ribotypes (VPI 630 or RT012 and VPI-10463 or RT078) used for challenge studies, showing comparable colony densities upon 1E4 dilution. *C. difficile* VPI-10463 was selected as the challenge strain for further efficacy experiments.
**(C)** Survival analysis of C57BL/6 mice challenged with different spore doses of RT078 shows that $10^5$ spores/mouse induce consistent and fulminant disease with >80% mortality, while $\leq 10^4$ spores produce a less severe phenotype allowing observation of recovery (n = 7).
**(D)** Body weight change over time in mice challenged with different spore doses confirms increased

disease severity with higher inocula (n = 7).

**(E)** Body weight progression comparing $VHH_4$ (B)-treated mice to Mock (B) controls shows significantly improved body weight on days 2 to 5 post-challenge in Exp 2 (n = 11, $p < 0.05$).

**(F)** Cox proportional hazards analysis reveals that $VHH_4$ treatment significantly reduces mortality compared to untreated controls (Hazard Ratio Exp 1 & Exp 2: 0.09, $p = 0.027$).

**(G**) Daily feed intake of mice across treatment groups confirms daily consumption of oral VHH4 treatment and a reduced intake in the VHH4-mIgA Fc group during the acute infection phase comparable with Untreated mice (n = 2 for Exp 1 and n = 3 for Exp 2).
